# Two for tau: Automated model discovery reveals two-stage tau aggregation dynamics in Alzheimer’s disease

**DOI:** 10.1101/2024.07.15.603581

**Authors:** Charles A. Stockman, Alain Goriely, Ellen Kuhl, and the Alzheimer’s Disease Neuroimaging Initiative

## Abstract

Alzheimer’s disease is a neurodegenerative disorder characterized by the presence of amyloid-*β* plaques and the accumulation of misfolded tau proteins and neurofibrillary tangles in the brain. A thorough understanding of the local accumulation of tau is critical to develop effective therapeutic strategies. Tau pathology has traditionally been described using reaction-diffusion models, which succeed in capturing the global spread, but fail to accurately describe the local aggregation dynamics. Current mathematical models enforce a single-peak behavior in tau aggregation, which does not align well with clinical observations. Here we identify a more accurate description of tau aggregation that reflects the complex patterns observed in patients. We propose an innovative approach that uses constitutive neural networks to autonomously discover bell-shaped aggregation functions with multiple peaks from clinical positron emission tomography (PET) data of misfolded tau protein. Our method reveals previously overlooked two-stage aggregation dynamics by uncovering a two-term ordinary differential equation that links the local accumulation rate to the tau concentration. When trained on data from amyloid-*β* positive and negative subjects, the neural network clearly distinguishes between both groups and uncovers a more subtle relationship between amyloid-*β* and tau than previously postulated. In line with the amyloid-tau dual pathway hypothesis, our results show that the presence of toxic amyloid-*β* influences the accumulation of tau, particularly in the earlier disease stages. We expect that our approach to autonomously discover the accumulation dynamics of pathological proteins will improve simulations of tau dynamics in Alzheimer’s disease and provide new insights into disease progression.

## 1. Introduction

Alzheimer’s disease affects the human brain years if not decades before the first symptoms of neurodegeneration emerge. Extracellular amyloid-*β* plaques and hyperphosphorylated tau protein play a major role in the pathology of neurodegeneration [37]. A comprehensive understanding of their contributions, their interplay, and their timeline of progression is crucial for the development of targeted treatments for Alzheimer’s disease.

Increasing evidence suggests that pathological proteins in neurodegenerative diseases spread in a prion-like manner, indicating that misfolded proteins can act as infectious agents that induce the misfolding and aggregation of healthy proteins, thereby transmitting the disease [8, 13, 39, 14, 25, 26, 36]. Mathematical modeling of misfolded protein aggregation, coagulation, and spreading hold promise to provide insight into the complex pathology of Alzheimer’s disease [11, 12, 9]. Integrated into machine learning, these models can successfully reproduce real life imaging data, make personalized assessments of disease progression [7, 32], and eventually correlate the concentration of misfolded protein to brain atrophy [31]. The mathematical framework to do this is rooted in reaction-diffusion-type models, including the single-field Fisher-Kolmogorov model [10, 16], the two-field Heterodimer model [29], and the multi-field Smoluchowski model [34]. A recent study focused on discovering the mathematical reaction-diffusion equation itself by combining physics-informed neural networks and symbolic regression [42].

All these models have one thing in common: They combine a linear diffusion and a nonlinear reaction term. The diffusion or spreading of misfolded tau protein and neurofibrillary tau tangles across the human brain follows a stereotypical distribution pattern commonly known as Braak stages [37, 41]. Braak stages I-II are characterized by the presence of misfolded tau in the transenthorinal layer, followed by Braak stages III-IV affecting the limbic system, and lastly Braak stages V-VI destructing virtually all isocortical association areas [4]. We can model this characteristic spreading of tau across the brain using weighted network models of the brain’s connectome, e.g., from the Budapest Reference Connectome [35] or the Human Connectome Project [23]. At the same time, the concrete aggregation mechanisms that drive the accumulation of misfolded tau protein remain poorly understood. Recent studies suggest that, from Braak stage III onward, local replication, rather than spreading between different brain regions, is the dominant mechanism that controls the overall rate of tau accumulation [24].

In this study, we neglect the diffusion of misfolded protein and focus exclusively on discovering the aggregation term for tau protein misfolding in the human brain using longitudinal real-life data. We design a constitutive neural network [19] to discover a bell-shaped function with multiple peaks that best describes clinical positron emission tomography (PET) data of misfolded tau protein. We train the network on publicly available imaging data from the Alzheimer’s Disease Neuroimaging Initiative [2], using longitudinal tau PET data of 401 subjects, each with two to six consecutive scans. We also study the influence of amyloid-*β* on the aggregation of tau, and discover two distinct equations for amyloid-*β* positive and negative subjects. Furthermore, we tailor the neural network to the specific regions of the Braak stages.

## 2. Methods

### 2.1 Image Data

We use longitudinal image data from the Alzheimer’s Disease Neuroimaging Initiative (ADNI). This database provides publicly available data and focuses on the early detection and tracking of Alzheimer’s disease [2]. From this dataset, we select a subset of 401 subjects that contain the standardized value uptake ratios (SUVRs) that measure the tau AV1451-PET and amyloid AV45-PET uptakes for 109 FreeSurfer-defined regions of the brain. For simplicity, we refer to these data sets as tau data and amyloid-*β* data throughout the remainder of this work. Each of the 401 subjects has two to six consecutive scans; 226 subjects are amyloid-*β* positive, 175 are amyloid-*β* negative.

#### Misfolded tau data

The processing of the tau AV1451PET data by ADNI follows standard protocols [3]. Briefly, the PET images for each subject are co-registered to the corresponding high-resolution T1-weighted magnetic resonance images closest in time to the relevant tau scan, and segmented into the FreeSurfer-defined regions of interest. For each region of interest, a standardized uptake value ratio *c*^img^ is calculated using the inferior cerebellum as the reference region. We include all right, left, and volume-weighted bilateral FreeSurfer-defined regions of interest and exclude the hippocampal region, which is often contaminated by offtarget binding. We group all composite regions of interest into the six Braak regions [4], see Figure 1. Since the Braak II region consists only of the hippocampus, we exclude this region in further analysis. We calculate two different variables. First, we normalize all data points in the unit interval, 0 ≤ *c*_all_ ≤ 1. We call these values the *whole brain-normalized concentration c*_all_. This variable will facilitate direct comparison between different regions of the brain.

**Figure 1:**
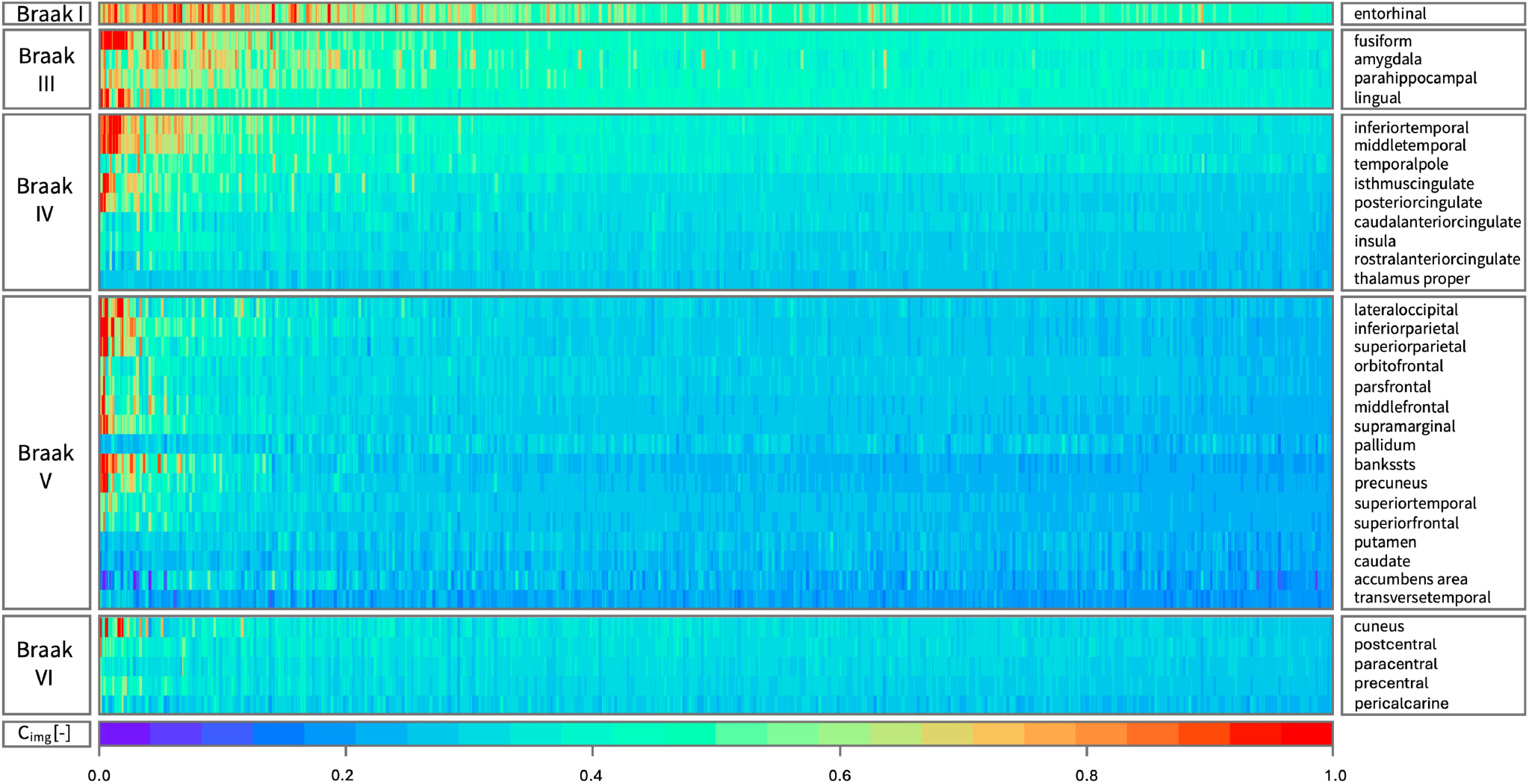
Longitudinal tau positron emission tomography concentration in Braak regions. Standardized uptake value ratios from positron emission tomography scans of 401 subjects with two to six annual scans in the 34 regions of the six Braak stages. The hippocampus, comprising the Braak II region, is excluded due to off-target binding contamination. Regions on the vertical axis are grouped per Braak region and sorted by mean tau load, from top to bottom. Subjects on the horizontal axis are sorted by mean tau load across all Braak regions and visits, from left to right. Each block of columns represents data for one subject. Within each block, each subcolumn represents data from one annual scan.

Second, we define the *region-normalized concentrations c*_roi_. We hypothesize that these values capture regional variations in production dynamics, while accounting for a similar aggregation mechanism. Per region of interest, we calculate the regional baseline value *c*_0_, and carrying capacities *c*_∞_, and normalize *c*_roi_ between *c*_0_ and *c*_∞_ [7, 37]. The former represents the tau concentration in the healthy state, the latter is the tau concentration in the late-stage Alzheimer’s disease state. We derive *c*_0_ and *c*_∞_ using a two-component Gaussian mixture model individually for each region of interest [37], resulting in Figure 5A. Lastly, we calculate the tau accumulation rate, or rate of change of the regionnormalized concentration *ċ* _roi_ using a forward Euler time integration,

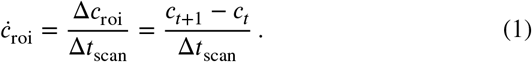

We divide the change in region-specific concentration between two consecutive scans Δ*c*_roi_ by the time between two scans Δ*t*_scan_ and seek to discover the aggregation equation that best describes the change in the misfolded protein concentration.

#### Amyloid beta data

We follow a similar workflow for the amyloid-*β* positron emission tomography data. The algorithm determines the amyloid SUVR values by coregistering amyloid AV45and NAV-PET scans with structural T1-weighted magnetic resonance images for each FreeSurfer Desikan-Killiany region (left, right, and volume-weighted bilateral), the cortical summary region (consisting of the frontal, anterior/posterior cingulate, lateral parietal and lateral temporal regions), and the five reference regions (cerebellar grey matter, whole cerebellum, brainstem, eroded subcortical white matter, and a composite reference region). In addition to the exact amyloid-*β* SUVR in each region of interest, the database provides the amyloid status for each subject [30]. We calculate the heterogeneous regional baseline values and carrying capacities to account for the spatio-temporal distribution of the protein concentration [40], and distinguish between the whole brain-normalized *c*_all_ and region-normalized *c*_roi_ concentrations.

### 2.2 Constitutive neural network

We use a constitutive neural network [19] to discover the fundamental relationship between the tau concentration and the tau accumulation rate, see Figure 2.

**Figure 2:**
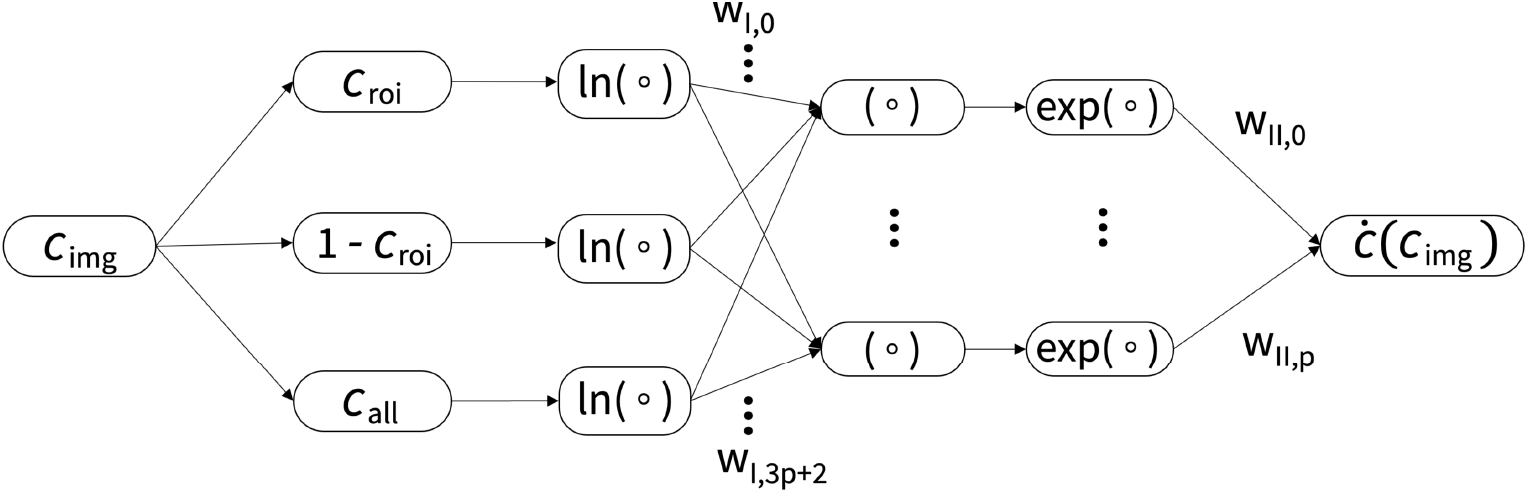
Constitutive neural network for protein misfolding.. The network represents the p-term aggregation function capable of describing the accumulation rate of misfolded tau in each region of the brain. It takes the whole brain-normalized concentration as input and computes the three concentrations, *c*_roi_, (1 − *c*_roi_), *c*_all_. From them, it calculates the powers for each concentration and multiplies the variables using a logarithmic-exponential transformation. Each term is weighted and added to the accumulation rate ċ (*c*) as output.

#### Discovering the fundamental aggregation equation

As our understanding of the underlying molecular and cellular mechanisms of tau aggregation remains incomplete, we propose a strategy to autonomously discover the equation that governs the aggregation, production, and accumulation of misfolded tau in each region of the brain, purely based on real-life clinical data [17, 21, 28]. We first convert the imaging data *C*_img_ into the region-normalized and whole brain-normalized concentrations,

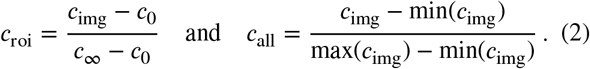

We then offer the discovery the following function, a sum of *p* individual contributions of three terms each,

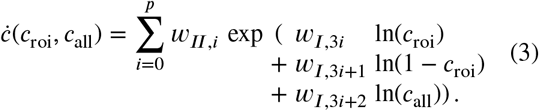

that we can convert into *p* products of three concentration terms,

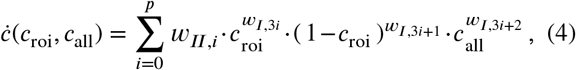

In our network equations (3) and (4), we neglect transport between regions and only focus on the local accumulation of misfolded protein. Figure 2 illustrates the constitutive neural network that takes the tau concentration *c*_img_ as input and computes three concentration terms. The first two terms, *c*_roi_ and (1 − *c*_roi_), are the region-normalized concentrations relative to the baseline value *c*_0_ and carrying capacity *C*_∞_ of the specific region. This scaling accounts for spatial inhomogeneities across the brain [7], while assuming a similar aggregation mechanism across all brain regions. Both terms ensure that the discovered function will go to zero at *c*_0_ and *c*_∞_. The third term, *c*_all_, is the whole brain-normalized concentration that captures the potential influence of the absolute concentration of misfolded protein. The network weights *w*_*I*_ represent the powers of each term, as previous mathematical aggregation models often involve powers of the concentration [11, 38]. Since our algorithm sums up the layers of the neural network sequentially [15], we first apply a logarithmic transformation to each variable, then calculate the weighted sum, and finally apply an exponential transformation, while constraining the weights to always remain non-negative, ***W*** ≥ 0. Lastly, we sum up the resulting *w*_*II*_ weighted product *p* times. The weights of the first layer *w*_*I*_ are always unit-less [-], while the weights of the second layer *w*_*II*_ have the unit of time [-/year] and allow for a clear physical interpretation.

This network generates *p* activation functions as output, which take the shape of *p* bell shaped functions. We can interpret these bell-shaped functions as derivatives of a modified sigmoid function. Essentially, by combining multiple activation functions of this kind, we enable the constitutive neural network to discover a bell-shaped function with multiple peaks. When integrating this multiple-peak function, we derive a monotonically increasing sigmoid function that consists of *p* segments.

During training, our constitutive neural network learns the network weights ***W*** = {*w*_*I*,1_, …, *w*_*I*,3*p*+2_, *w*_*II*,1_, …, *w*_*II,p*_}, by minimizing a loss function *L* [22]. This loss function is the mean-squared error, or the *L*_2_-norm of the difference between the model, *ċ* (*c*_img_), and the training data, 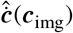, across all regions divided by the number of training points *n*_train_,

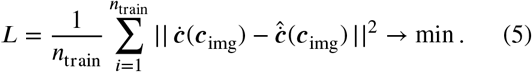

We train the network by minimizing the loss function (5) and learn the network parameters ***W*** using the ADAM optimizer, a robust adaptive algorithm for gradient-based first-order optimization, and constrain the weights to always remain non-negative, ***W*** ≥ 0 [20, 27].

#### Special cases

Our constitutive neural network can recover classical growth equations as special cases, for example, the Fisher-Kolmogorov-Petrovsky-Piskunov equation. Currently, this equation is mainly used to describe the relationship between tau and the accumulation rate of tau. We recover the local Fisher-Kolmogorov-Petrovsky-Piskunov equation [7] by looking at only one product term, *p* = 0, in equation (3), setting the region-normalized network weights to one, *w*_*I*,0_ = 1, *w*_*I*,1_ = 1, the brain-normalized network weight to zero, *w*_*I*,2_ = 0, and selecting appropriate values for *w*_*II*,0_. This leads to the following equation,

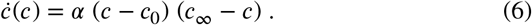

Naturally, our constitutive neural network includes different variations of the Kolmogorov-Petrovsky-Piskunov equation such as the Newell-Whitehead-Segel and Zeldovich-Frank-Kamenetskii equations [16, 42], see Table 1. To probe the performance of our network on synthetic data, we set the accumulation rate *ċ* (*c*) equal to the general form of their distributions, right column of Table 1, and add a small perturbation *ε* of 0.01 to simulate possible noise in the dataset. Furthermore, we calibrate our neural network on simple mathematical functions such as the constant, linear, and exponential function.

**Table 1:** Special cases of the constitutive neural network. The constant, linear, and exponential functions, as well as the Fisher-Kolmogorov-Petrovsky-Piskunov, Newell-Whitehead-Segel, and Zeldovich-Frank-Kamenetskii equations are special cases of our constitutive neural network in Figure 2. The right column provides their individual accumulation rates *ċ*(*c*).

| Equation | General form | Color |
| --- | --- | --- |
| Constant function | $\alpha$ | orange |
| Linear function | $\alpha c$ | purple |
| Exponential function | $\alpha e^{\beta(c-1)}$ | yellow |
| Fisher-Kolmogorov-Petrovsky-Piskunov | $\alpha c(1 - c)$ | red |
| Newell-Whitehead-Segel | $\alpha c(1 - c^q)$ | blue |
| Zeldovich-Frank-Kamenetskii | $\alpha c(1 - c)e^{\beta(c-1)}$ | green |

#### Prediction

After training the network, we use a modified forward Euler method to solve the differential equation for the tau accumulation rate as a function of the tau concentration. In this process, we fix the starting standardized value uptake ratio, and solve for time steps of 0.25 years. The resulting curve is smooth and continuous, with the possibility of segments with different slopes.

## 3. Results

### 3.1 Network calibration on synthetic data

First, we calibrate the network on synthetic data. We use artificially generated data from the constant, linear, and exponential function, as well as the Fisher-KolmogorovPetrovsky-Piskunov, Newell-Whitehead-Segel, and Zeldovich-Frank-Kamenetskii equations in Table 1. After training, we observe that our network consistently re-discovers the desired functions in less than 50 epochs. Figure 3 shows the discovered functions for the synthetic data of the Fisher-Kolmogorov-Petrovsky-Piskunov (blue), NewellWhitehead-Segel (red), and Zeldovich-Frank-Kamenetskii (green) models. Notably, the mean-squared error is consistently smaller than 1.45 · 10_−4_.

**Figure 3:**
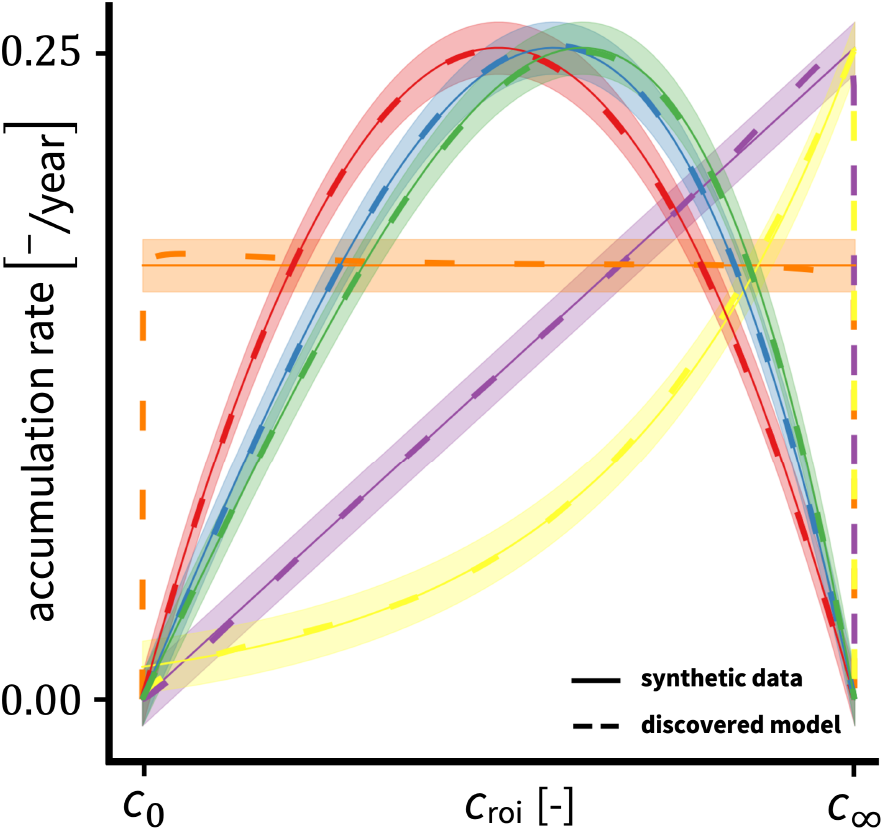
Model calibration on synthetic data. Constitutive neural network trained on synthetic data of three standard mathematical equations and three versions of the KolmogorovPetrovsky-Piskunov equation. Each case uses 10,000 synthetically generated data points with a small perturbation *ε*. The solid line shows the synthetic data; the dashed line shows our discovered model. The three standard mathematical equations include the constant function in orange, the linear function in purple, and the exponential function in yellow. The three versions of the Kolmogorov-Petrovsky-Piskunov equation include the Fisher-Kolmogorov-Petrovsky-Piskunov in blue, the Newell-Whitehead-Segel equation in red, and the ZeldovichFrank-Kamenetskii equation in green. For each distribution, the mean-squared error is smaller than 1.45 · 10^−4^.

### 3.2 Network training on combined subject data

Second, we train our network on real tau concentration data. Initially, we combine all *n*_train,all_ = 19, 680 data points from all subjects and all regions of interest of the brain. We initialize the weights of the neural network using the same values as during the calibration step.

We train the model (2) with *p* = 2 activation functions on the tau concentration data of all brain regions and subjects combined. We do not distinguish between amyloid*β* status or take into account age, sex, genotype, or the clinical diagnosis of the subject. The model fits the data in less than 250 epochs, and takes less than a minute to train. Importantly, the ADNI dataset contains image data, and most likely contains noise attributed to the affinity of the tau AV1451-PET tracer, the sensitivity of the imaging technique, and the image processing. To account for this noise, we use bootstrapping to estimate the variability of the learned function. We resample the dataset 1,000 times with replacement. All samples are independent of one another, and together used to estimate model uncertainty.

Figure 4 illustrates the result of the bootstrapping. It shows the tau accumulation rate versus the region-normalized tau for *n*_data,*Aβ*+_ = 41, 442 data points of the amyloid-*β* positive subjects in blue and for *n*_data,*Aβ*_ = 28, 466 data points of the amyloid-*β* negative subjects in orange. The black line indicates the mean and the grey shaded area is the confidence interval created by the bootstrapping. For training, we use all *n*_train,all_ = 19, 680 data points, and only include data for which the brain-normalized tau value lies between *c*_0_ and *c*_∞_, and for which the accumulation rate is non-negative, [inerq] ≥ 0. After training, we see that the aggregation function consistently takes the form of a bell-shaped function with two distinct peaks.

**Figure 4:**
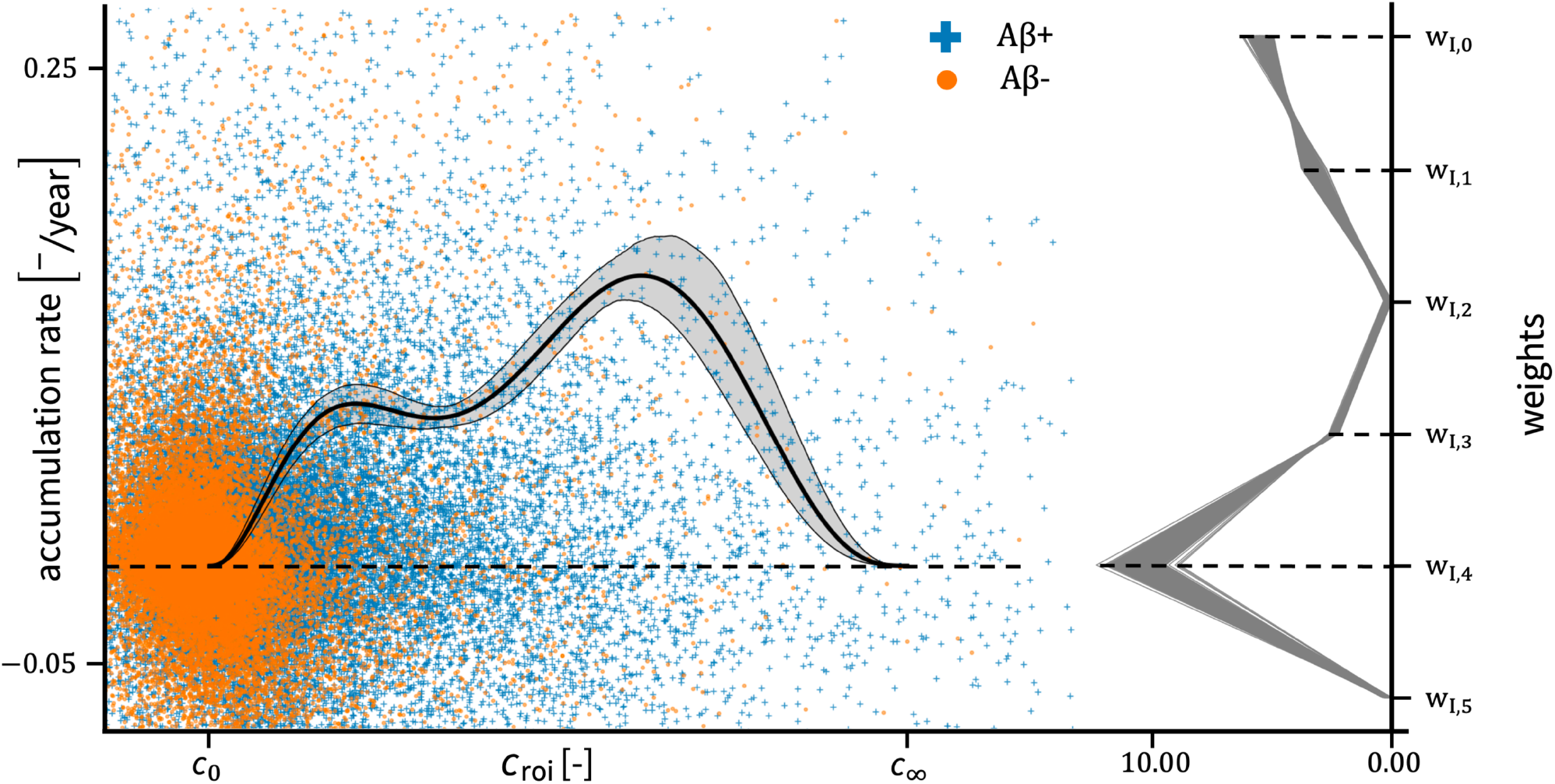
Accumulation rate as a function of region-normalized tau PET standardized uptake value ratio (SUVR). Scatterplot of annual accumulation rate of tau PET standardized uptake value ratios (SUVR) vs region-normalized tau PET (SUVR) for all subjects (*n*_all_ = 401). The subjects are divided into an amyloid-*β* positive group (+, *n*_*Aβ*+_ = 226) and an amyloid-*β* negative group (·, *n*_*Aβ*−_ = 175). The estimated function for the accumulation rate as a function of the region-normalized tau PET is shown with a 95 percent confidence interval (left), together with the trained network weights (right). The uncertainty in the accumulation rate is calculated by bootstrapping the constitutive neural network 1000 times, with 250 epochs per sample. The network fits the data robustly with a double bell-shaped function, accounting for two peaks or acceleration stages in the accumulation rate. Furthermore, the brain-normalized network weights *w*_*I*,2_ and *w*_*I*,5_ train consistently to zero, showing that the network prioritizes a dependence on the region-normalized concentration *c*_roi_ over the whole brain-normalized concentration *c*_all_.

The right graph in Figure 4 shows that the weights *w*_*I*,2_ and *w*_*I*,5_ that correspond to the brain-normalized concentration in equation (3) consistently train to zero. As the weight, and thus the power, go to zero, the contribution of the brainnormalized concentration *c*_all_ becomes negligible and the discovered function simplifies to a two-term function,

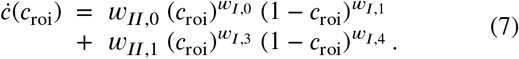

This function is completely defined within the interval [ *c*_0_, *c*_∞_ ]. Now, we translate the discovered function (7) back to the original interval for each region of interest, as denoted by Figure 5A. For this, we simply use the inverse calculation of the accumulation rate (1) and the definition of the region-normalized concentrations (3). The translation back to the relevant region results in Figure 5B. The regions are again sorted per Braak stage, and the discovered function (7) is plotted between the baseline value and carrying capacity of that relevant region. Importantly, we use the non-normalized values for the tau concentration and accumulation rate, allowing for better interpretation of the real values obtained by PET imaging. We can see that regions with a larger interval [ *c*_0_, *c*_∞_ ] have higher values for the accumulation rate.

**Figure 5:**
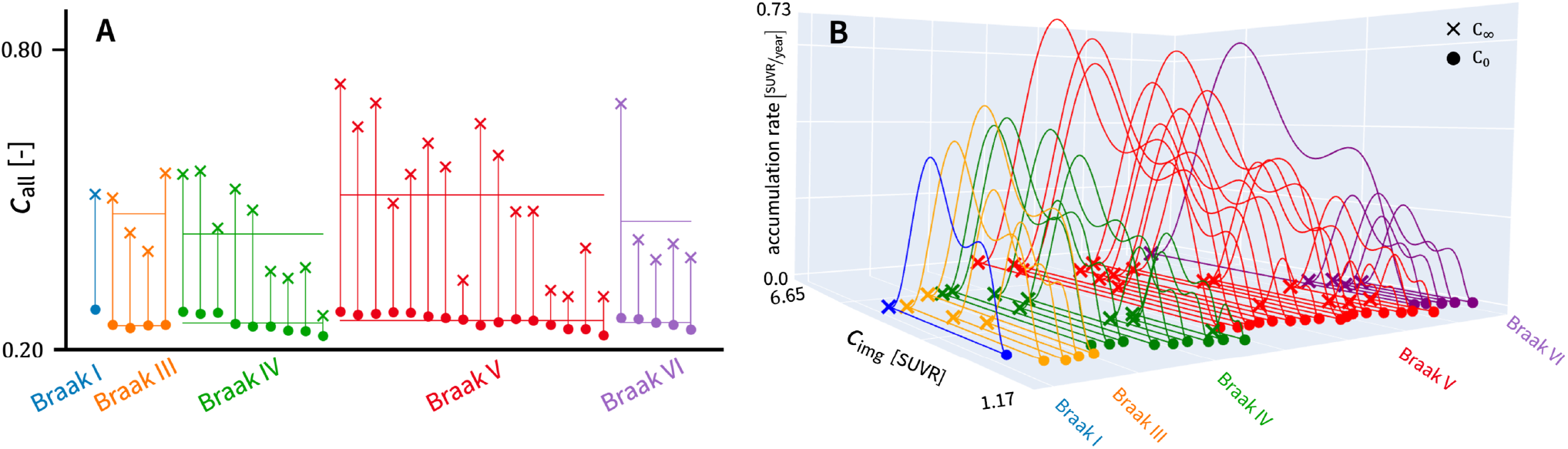
Regional baseline values (·) and carrying capacities (×) for brain-normalized tau and their accumulation rate. (a) For each region of interest of the brain, we fit a two-component Gaussian mixture model to tau data to derive the baseline value *c*_0_ (·) and the carrying capacity *c*_∞_ (×). We sort all regions per Braak stage similar to Figure 1 and calculate the mean per Braak stage for both parameters (_) for comparison. The tau baseline values for all regions are centered around *c*_all_ = 0.256, while the carrying capacities vary between 0.268 and 0.874. (b) Translation of the accumulation rate in Figure 4 to each region of interest as denoted in (a). Here, the non-normalized values for the tau concentration and accumulation rate are used. The magnitude of the accumulation rate depends on the size of the [*c*_0_, *c*_∞_]-interval.

### 3.3 Distinction by amyloid-β status

Third, we split the cohort between the *n*_train,*Aβ*+_ = 14, 808 amyloid-*β* positive subjects and the *n*_train,*Aβ*−_ = 4, 872 amyloid-*β* negative subjects. We use the weights provided by the calibration step to initialize the weights of the neural networks. We resample both datasets with replacement and create 1,000 bootstrap samples for both amyloid groups.

Figure 6A shows the comparison between the aggregation function for the amyloid-*β* positive group (orange) and the aggregation function for the amyloid-*β* negative group (blue), alongside their respective confidence intervals. Notably, the 95 percent uncertainty interval for the amyloid*β* negative group is wider than for the amyloid-*β* positive group. This can be attributed to the quantity of data points available for each group. Recall that we only take into account the concentrations for each region between the baseline value and the carrying capacity. This results in 14,808 data points for the amyloid-*β* positive group and only 4,872 data points for the amyloid-*β* negative group. We have thus more than three times the amount of data points for the amyloid-*β* positive group, resulting in a slimmer uncertainty interval. Moreover, the uncertainty in the accumulation rate of tau is smaller for a lower value of the region-normalized tau concentration *c*_roi_. This is to be expected, as the density of the data points for each group in Figure 4 is higher for lower values of *c*_roi_. Strikingly, there is a significant difference between both curves. The accumulation rate for the amyloid-*β* positive group clearly displays two distinct peaks. Initially, it increases faster than for the negative group, as evidenced by the first peak at *c*_roi_ = 0.203, and second peak at *c*_roi_ = 0.663. In contrast, the accumulation rate for the amyloid-*β* negative group shows only one local maximum, at *c*_roi_ = 0.577. The two groups thus exhibit a different behavior during the first stage of the disease, with a higher accumulation rate for the amyloid-*β* positive group. During later stages, after the intersection of the two curves at *ċ* (*c*_roi_ = 0.345) = 0.076, both groups behave quite similarly. However, the amyloid-*β* positive group already achieves the same value for *ċ* _*Aβ*+_ for a region-normalized concentration of *c*_roi_ = 0.148. Thus, in the case of the amyloid-*β* negative group, the region-normalized concentration needs to be substantially higher, i.e., more than twice as high, to reach the same accumulation rate as the amyloid-*β* positive group.

**Figure 6:**
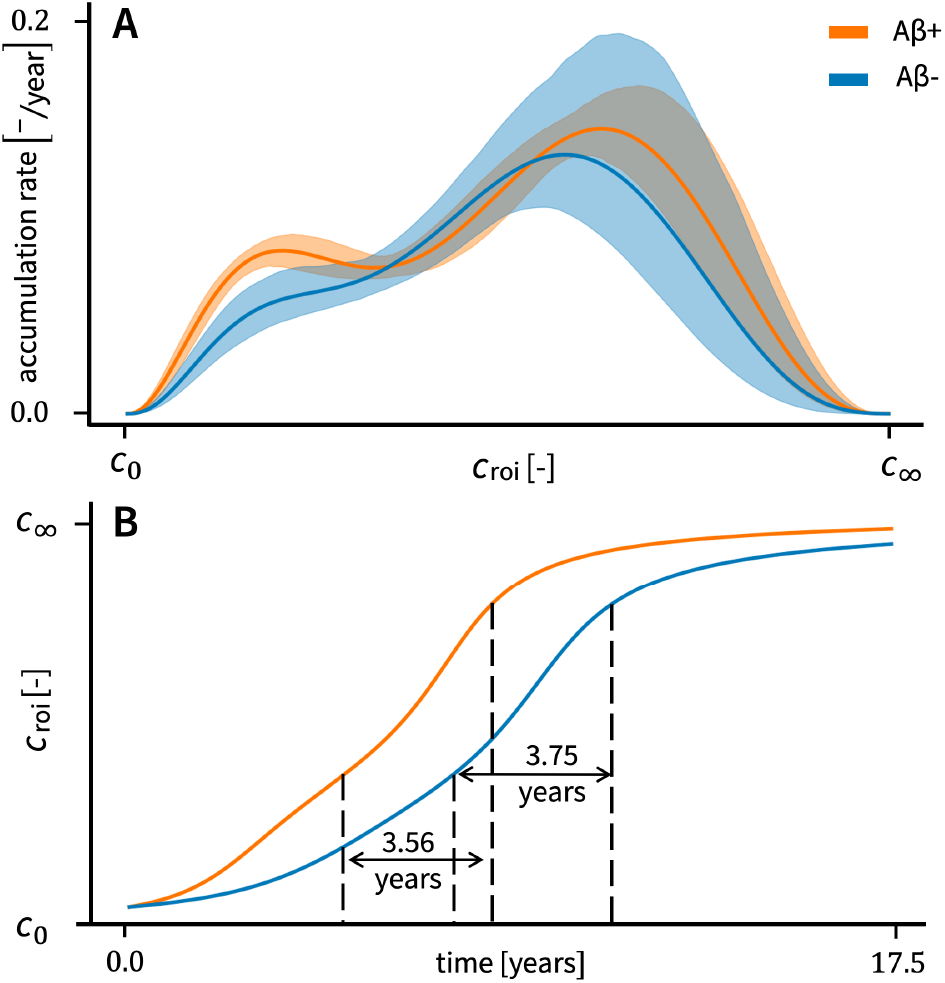
Prediction of the region-normalized tau PET concentration based on the modeled accumulation rate for amyloid-*β* positive (orange) and negative (blue) status. (a) The estimated function for the accumulation rate as a function of the region-normalized tau PET for amyloid-*β* positive (orange) and negative (blue) status, together with their 95 percent confidence intervals. The confidence interval for the amyloid-*β* negative group (*n*_train,*Aβ*−_ = 4872) is notably wider than the confidence interval for the amyloid-*β* positive group (*n*_train,*Aβ*+_ = 14808). (b) Integral with respect to time of the curves in (a) using a modified forward Euler method. In the case of the amyloid-*β* negative group, our model predicts that it takes 7.438 years to reach a concentration *c*_roi_ of 0.345, and subsequent 3.75 years to reach *c*_roi_ = 0.75. For the amyloid-*β* positive group in contrast, it takes only 4.875 years to reach a concentration *c*_roi_ of 0.345, and subsequent 3.56 years to reach *c*_roi_ = 0.75.

The effect of amyloid-*β* on the accumulation rate of tau becomes more intuitive when we integrate equation 3 with respect to time, see Figure 6B. At time *t* = 0, we assign the starting concentration of *c*_roi_ to 0.05. The time it takes to achieve a concentration *c*_roi_ of 0.345 is 4.875 years for the amyloid-*β* positive group, and 7.438 years for the amyloid-*β* negative group. In contrast, it takes only 3.56 years for the A*β*+ group and 3.75 years for the A*β*group to increase another 40% in concentration.

### 3.4 Distinction by Braak stages

Lastly, we train the network on the regions of the different Braak stages individually, for Braak stage I with *n*_*I,Aβ*−_ = 86 and *n*_*I,Aβ*+_ = 217, for stage III with *n*_*III,Aβ*−_ = 99 and *n*_*III,Aβ*+_ = 255, for stage IV with *n*_*IV, Aβ*−_ = 318 and *n*_*IV, Aβ*+_ = 749, for stage V with *n*_*V, Aβ*−_ = 524 and *n*_*V, Aβ*+_ = 1475, and for stage VI with *n*_*VI,Aβ*−_ = 64 and *n*_*VI,Aβ*+_ = 113. We initialize the weights of the neural networks using the same values as during the calibration step. We resample both datasets with replacement and create 500 bootstrap samples for each Braak stage.

Figure 7 shows the comparison between the aggregation function for the different Braak stages for the amyloid-*β* negative group (left column) and positive group (right column). For both amyloid-*β* status, we visualize the aggregation curves with their confidence intervals (upper row), and subsequently integrate the curves to obtain the prediction of the region-normalized tau concentration (lower row). We color-code the curves by Braak stage with stage I (blue), stage II (orange), stage IV (green), stage V (red), and stage VI (purple). Notably, the confidence intervals for each of the cases are wider than in Figures 4 and 6. This is a result of the lower amount of training points per Braak stage, which contains only about a sixth of data points than the data split solely by amyloid-*β* status. Another effect of the small amount of data points per training region is that the confidence intervals are less smooth and exhibit more artifacts compared to the previous cases. This results in a greater impact of randomly not choosing certain data points during bootstrapping.

**Figure 7:**
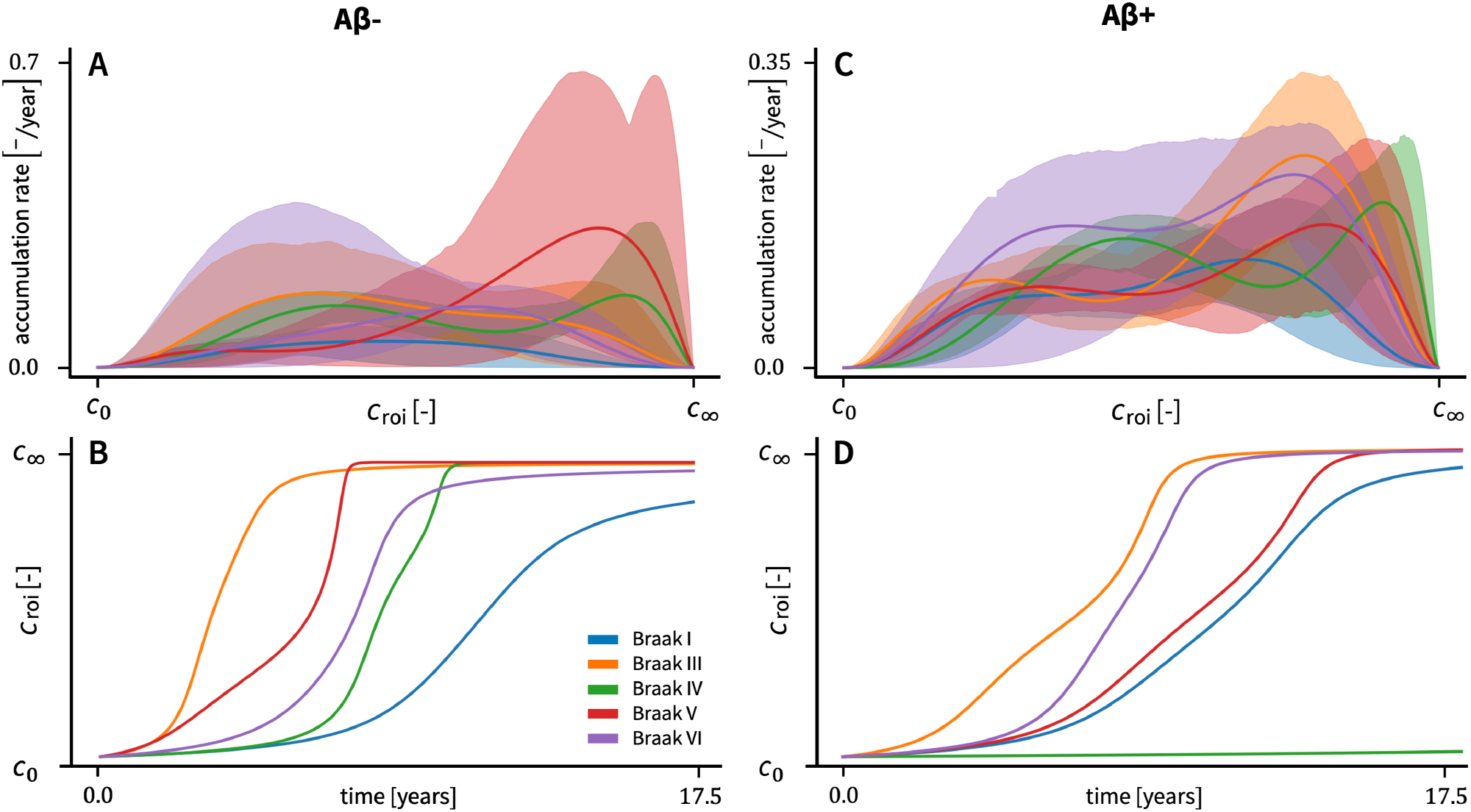
Prediction of the region-normalized tau PET concentration based on the modeled accumulation rate for the different Braak stages, for amyloid-*β* negative (left) and positive (right) status. The estimated function for the accumulation rate as a function of the region-normalized tau PET for Braak stage I (blue), stage II (orange), stage IV (green), stage V (red), and stage VI (purple), together with their 95 percent confidence intervals for (a) the amyloid-*β* negative group and (c) the amyloid-*β* positive group. There is a big overlap between the confidence intervals of the different Braak stages, and the confidence intervals are significantly wider than in the previous cases. Using a modified forward Euler method, the prediction of the accumulation of the protein tau for (b) the amyloid-*β* negative group and (d) the amyloid-*β* positive group is given. In (b), we see the fastest increase in the concentration of tau happening for Braak II, followed by Braak V, Braak VI, Braak IV, and finally Braak I. In (d), we see the fastest increase in the concentration of tau happening for Braak II, followed by Braak VI, Braak V, Braak I, and finally Braak IV. The concentration in Braak IV only starts to increase after a period of 40 years, due to the low values for the accumulation rate for values closer to the baseline value in (c).

When comparing the Braak-specific curves for both amyloid-*β* status, we expect to observe the same behavior as in Figure 6, that is, the amyloid-*β* positive curve is above the amyloid-*β* negative curve for lower values of the regionnormalized concentration *c*_roi_, and similar curves for higher values. Strikingly, we only see this behavior in the curves for Braak stages I and VI. For Braak stage III, the amyloid-*β* negative curve is higher for lower values, with the positive curve becoming larger around a value of *c*_roi_ = 0.6. In Braak stage IV, the curves for the accumulation rate are quite similar, with the amyloid-*β* negative curve slightly shifted to lower values of *c*_roi_. For Braak stage V, the amyloid-*β* negative curve is smaller for lower values as expected, but it becomes significantly larger for higher values of *c*_roi_.

This unexpected behavior becomes more visible when integrating the mean of the Braak-specific curves with respect to time, resulting in Figure 7B and 7D. Generally, the concentration of tau for the amyloid-*β* negative status increases faster than for the positive status, apart from Braak stages I and VI. Even more unexpected is the extremely late increase in tau concentration for the amyloid-*β* positive Braak stage IV.

## 4. Discussion

In this study, we propose a new way of autonomously discovering the accumulation of misfolded tau protein in different regions of the brain, purely derived from clinical imaging data. Therefore, we designed a constitutive neural network [19] capable of learning a function with multiple peaks. These multiple peaks could potentially represent multiple accelerations and decelerations in the progression of the accumulation of tau, and thus lead to a better understanding of the progression of Alzheimer’s disease in general. We calibrated the model using longitudinal tau PET data of 401 subjects. Encouraged by the fast training time of the network, we used bootstrapping to estimate the uncertainty of our model and calculated 95 percent confidence intervals.

We ensured that the constitutive neural network is capable of mimicking the shape of standard simple mathematical equations as well as known growth equations currently used for the longitudinal modeling of tau PET data in Table 1. The ability of our neural network to reproduce these equations suggests that, if the accumulation of the protein tau mostly follows a single bell-shaped function, our neural network will be able to accurately capture this behavior. However, when trained on real tau PET data, we see that the constitutive neural network robustly trains to a bell-shaped function with two peaks, see Figure 4. This suggests that there are two distinct stages in the progression of Alzheimer’s disease: a first stage at the beginning of the disease when the concentration of tau increases quite rapidly, followed by second stage with a steady accumulation rate. The transition between both stages is characterized by a notable increase in the accumulation rate.

The introduction of the region-specific baseline values and carrying capacities for tau plays a major role in the discovery of the two-stage behavior. We defined these values based on all the data points of tau for a certain region. To compare their relative importance, we calculated the whole brain-normalized and region-normalized tau concentrations, and integrated both in the constitutive neural network. During training, it became obvious that our neural network prioritizes the region-normalized over brain-normalized tau concentrations. More specifically, the network weights associated with the brain-normalized tau concentration completely trains to zero. This suggests that the accumulation rate of tau only depends on the region-normalized concentrations *c*_roi_, and not on the brain-normalized concentration *c*_all_. For our discovered model, we conclude that the baseline value and carrying capacity are indeed sufficient to represent accurately the process of tau accumulation. We can also interpret the interval between the baseline value and carrying capacity as the timeline of Alzheimer’s disease, within which all the aggregation dynamics of tau occur that contribute to neurodegeneration. Specifically, we can interpret the baseline value as the onset of the aggregation of tau, exhibiting the same characteristics as an unstable equilibrium. If the concentration in a specific region is smaller than this baseline value, tau will not increase. Once the concentration exceeds the baseline value, the tau concentration will keep growing until it reaches a plateau, the carrying capacity. The carrying capacity is the maximum concentration of misfolded tau that a region of the brain can contain.

However, the definition of the baseline value and carrying capacity also limits our automated model discovery. These two variables are derived from fitting a two-component Gaussian mixture model to the tau data of that specific region of the brain. These values depend on the imaging data, and they can slightly vary if we include or exclude data from the dataset. After finding these values, we enforce the discovered function to zero at both the baseline value and the carrying capacity, fixing the discovered function within this interval. A potential next step would be to include the discovery of the baseline value and carrying capacity in our neural network. This would enable the translation of the function in the concentration direction, and give the neural network more freedom in the discovery of the accumulation rate.

Once we have discovered the general function for the accumulation rate, we translate it back to the original regions of interest, resulting in Figure 5. Here, the notion of the regional baseline value and carrying capacity becomes more intuitive. We hypothesize that these variables represent the regional variations in production dynamics, while accounting for a similar aggregation mechanism. The regional variation is captured by the limits imposed by the baseline value and carrying capacity. This means that tau aggregation in the relevant region of the brain only occurs within the [ *c*_0_, *c*_∞_ ]interval. The similar aggregation mechanism is represented by the same general mathematical function found by our neural network. However, the magnitude of the function depends on the size of the [ *c*_0_, *c*_∞_ ]-interval. We can clearly see that the introduction of region-specific baseline values and carrying capacities allows us to capture variations across the different regions of the brain.

We also investigated the potential influence of amyloid-*β* on the accumulation of the protein tau. This is motivated by the amyloid-tau dual pathway hypothesis, which postulates that amyloid and tau conspire with one another to enhance each others’ toxicity [33, 36]. Here, our constitutive neural network was able to distinguish between the accumulation dynamics based on amyloid-*β* status. We found that the presence of toxic amyloid-*β* influences the accumulation rate of the protein tau. More specifically, amyloid-*β* increases the accumulation of the protein tau when the tau concentration is smaller than a certain threshold. Beyond that threshold, the presence of toxic amyloid-*β* no longer accelerates the accumulation of tau, and the tau accumulation rate becomes independent of amyloid status. Recent studies have shown that toxic amyloid-*β* plaques provide a microenvironment that promotes tau aggregation and propagation [6, 18]. This suggests that the presence of amyloid-*β* fosters the initial seeding and hyperphosphorylation of healthy tau. However, once a sufficient amount of healthy tau has been converted into its pathological form, it is mainly the presence of other toxic tau that drives the accumulation and recruitment of additional healthy tau protein.

While the distinction between amyloid status shows a clear difference in the accumulation of tau, an important next step is to include the specific values of the concentration of amyloid in the relevant region. This will help us to gain more computational insights into the interaction between amyloid-*β* and tau. It is important to note that the numerical results regarding the difference between tau accumulation for the amyloid-*β* positive and negative groups are based on the mean of the bootstrapped function in Figure 6A. In reality, the exact difference in time might differ from our inferred results. The important observation is that there is a significant difference between the accumulation rate for both groups, suggesting that the presence of toxic amyloid*β* should be included in future tau aggregation models as argued in [36, 5] and found necessary to explain changes in brain activity during the disease [1]

Finally, we looked at the individual Braak stages to discover the accumulation rate per Braak region. Yet, the outcome differed from our expected results. Per Braak stage, we expected to find a similar behavior for both amyloid groups as for the whole brain in Section 3.3, with a faster increasing concentration for the amyloid-*β* positive group than for the amyloid-*β* negative group. However, for the Braak stages III, IV, and V, we found the opposite behavior: the concentration for the amyloid-*β* negative group increases faster than for the amyloid-*β* positive group. In addition, we anticipated our neural network to replicate the Braak sequence, with an initial increase for Braak stage I, followed by Braak stages III to VI. In contrast, our model does not replicate this sequence. Importantly, our discovered functions represents the local aggregation dynamics of the protein tau, starting from the baseline value *c*_0_. Our study does not attempt to derive the exact timeline of regional tau activation. If this is the goal, we need to combine our model with a diffusion model of the brain that allows for initial seeding of misfolded tau protein until the baseline value is reached. We hypothesize that once the tau concentration exceeds this baseline value, it is dominantly the local aggregation that will control the overall rate of tau accumulation as also found in [7].

When we examine the training data and the discovered functions for each Braak stage in more detail, we can identify different families of discovered functions, corresponding to certain outliers in the training set. The bootstrapping randomly includes or excludes data points in the training set. We see that when points with extremely large values for the accumulation rate are included, our neural network is drawn to these points, trying to fit the data as accurately as possible. As an example, in the case of the amyloid-*β* negative Braak stage IV and V, there is a data point with a larger value of the tau concentration and a relatively high accumulation rate. This point forces the model discovery closer to the carrying capacity, resulting in a first family of discovered functions. This effect, combined with the constraint on the function to be zero at the carrying capacity, explains the sudden flattening near the carrying capacity observed in Figure 7B for both Braak stages. When this point is excluded from the training set, the discovered model give a second family of derived functions consistent with our expectations. We observe a similar multi-family discovery for other Braak stages. This variability indicates that the data set for the individual Braak stages either lacks sufficient information to train our network robustly or contains too many outliers.

In the future, tailoring this neural network to specific patient data will be necessary to discover accurate personalized models of Alzheimer’s disease pathology and progression. From these, the network for tau accumulation should be extended to include the discovery of brain atrophy and its relation to the local tau concentration. Together with the integration of the concrete concentration of amyloid in the network, this will help uncover the intricate relationship between tau, amyloid, and neurodegeneration. By directly linking the concentration of misfolded protein to brain atrophy, we can define a threshold for the region normalized-concentration at which neurodegeneration occurs.

## 5. Conclusion

Current reaction-diffusion equations used to model the accumulation of tau in Alzheimer’s disease have succeeded in capturing the spread of misfolded tau protein and neurofibrillary tangles, but fail to accurately characterize the local aggregation, production, and accumulation of tau. Here we present a novel strategy to autonomously discover tau accumulation functions with possibly multiple peaks. When applied to clinical PET data of misfolded tau protein, we discover physics-based models that capture the underlying dynamics linking local accumulation rate and local tau concentration. Surprisingly, we discover, robustly, a two-stage aggregation dynamics. This observation is in contrast with existing tau accumulation models that use mathematical expressions with a single peak to explain clinical data. The two-peak behavior of the accumulation rate translates into two distinct stages with an increased tau concentration.

When distinguishing between amyloid-*β* status, the network robustly discovers significantly different aggregation dynamics for the positive and negative groups. Consistent with the amyloid-tau dual pathway hypothesis, our analysis supports the enhancement of tau protein accumulation in the presence of amyloid, especially during the early stages of neurodegeneration. Our study provides new insights into the progression of Alzheimer’s disease and offers opportunities to tailor these insights into personalized timelines of disease progression.

## Acknowledgments

This research was supported by a B.A.E.F. master’s level research fellowship to Charles A. Stockman, and by the NSF CMMI Award 2320933 *Automated Model Discovery for Soft Matter* and the ERC Advanced Grant 101141626 *DISCOVER* to Ellen Kuhl.

Data collection and sharing for the Alzheimer’s Disease Neuroimaging Initiative (ADNI) is funded by the National Institute on Aging (National Institutes of Health Grant U19 AG024904). The grantee organization is the Northern California Institute for Research and Education. In the past, ADNI has also received funding from the National Institute of Biomedical Imaging and Bioengineering, the Canadian Institutes of Health Research, and private sector contributions through the Foundation for the National Institutes of Health (FNIH) including generous contributions from the following: AbbVie, Alzheimer’s Association; Alzheimer’s Drug Discovery Foundation; Araclon Biotech; BioClinica, Inc.; Biogen; Bristol-Myers Squibb Company; CereSpir, Inc.; Cogstate; Eisai Inc.; Elan Pharmaceuticals, Inc.; Eli Lilly and Company; EuroImmun; F. Hoffmann-La Roche Ltd and its affiliated company Genentech, Inc.; Fujirebio; GE Healthcare; IXICO Ltd.; Janssen Alzheimer Immunotherapy Research & Development, LLC.; Johnson & Johnson Pharmaceutical Research & Development LLC.; Lumosity; Lundbeck; Merck & Co., Inc.; Meso Scale Diagnostics, LLC.; NeuroRx Research; Neurotrack Technologies; Novartis Pharmaceuticals Corporation; Pfizer Inc.; Piramal Imaging; Servier; Takeda Pharmaceutical Company; and Transition Therapeutics.

